# Genome-wide identification and expression specificity analysis of the DNA methyltransferase gene family under adversity stresses in cotton

**DOI:** 10.1101/411652

**Authors:** Xiaomin Yang, Xuke Lu, Xiugui Chen, Delong Wang, Junjuan Wang, Shuai Wang, Lixue Guo, Chao Chen, Xiaoge Wang, Binglei Zhang, Mingge Han, Wuwei Ye

## Abstract

DNA methylation is an important epigenetic mode of genomic DNA modification that is an important part of maintaining epigenetic content and regulating gene expression. DNA methyltransferases (MTases) are the key enzymes in the process of DNA methylation. Thus far, there has been no systematic analysis the DNA MTases found in cotton. In this study, the whole genome of cotton C5-Mtase coding genes was identified and analyzed using a bioinformatics method based on information from the cotton genome. In this study, 51 DNA MTase genes were identified, of which 8 belonged to *G. raimondii* (group D), 9 belonged to *G. arboretum* L. (group A), 16 belonged to *G. hirsutum* L. (group AD_1_) and 18 belonged to *G. barbadebse* L. (group AD_2_). Systematic evolutionary analysis divided the 51 genes into four subfamilies, including 7 MET homologous proteins, 25 CMT homologous proteins, 14 DRM homologous proteins and 5 DNMT2 homologous proteins. Further studies showed that the DNA MTases in cotton were more phylogenetically conserved. The comparison of their protein domains showed that the C-terminal functional domain of the 51 proteins had six conserved motifs involved in methylation modification, indicating that the protein has a basic catalytic methylation function and the difference in the N-terminal regulatory domains of the 51 proteins divided the proteins into four classes, MET, CMT, DRM and DNMT2, in which DNMT2 lacks an N-terminal regulatory domain. Gene expression in cotton is not the same under different stress treatments. Different expression patterns of DNA MTases show the functional diversity of the cotton DNA methyltransferase gene family. VIGS silenced Gossypium hirsutum l. in the cotton seedling of DNMT2 family gene *GhDMT6*, after stress treatment the growth condition was better than the control. The distribution of DNA MTases varies among cotton species. Different DNA MTase family members have different genetic structures, and the expression level changes with different stresses, showing tissue specificity. Under salt and drought stress, *G. hirsutum* L. TM-1 increased the number of genes more than *G. raimondii* and *G. arboreum* L. *Shixiya* 1. The resistance of Gossypium hirsutum L.TM-1 to cold, drought and salt stress was increased after the plants were silenced with *GhDMT6* gene.

## Introduction

DNA methylation is the process of transferring a methyl (-CH_3_) group to a specific base of a DNA molecule and is catalyzed by DNA MTases, with S-adenosine methionine (SAM) as a methyl donor [1, 2]. DNA methylation widely occurs in the epigenetics of bacteria, plants, and animals and is involved in transposons [3-6], the suppression of gene silencing, genomic imprinting [7], X chromosome inactivation [8], cell differentiation [9], and embryo development [10]. However, DNA methylation is not immutable; under conditions of stress, the plant genome can overcome the limitation of genome instability through DNA methylation by rapid modification. This induces the expression of some genes associated with stress to maintain plant growth and development and evolutionary process [11-13]. Therefore, epigenetic modification precedes genomic evolution in response to adversity, and DNA methylation is considered the molecular response mechanism of plants in the face of adverse stress [5, 6, 11].

DNA methylation occurs mostly in CpGs at carbon 5 in cytosine (C5). It primarily occurs in symmetric sequence CGs but also occurs in CHG and CHH (H=A, C or T) sequences [14]. There are two DNA methylation methods in plants: maintenance methylation and denovo methylation [15]. Maintenance methylation refers to the methylation of a chain of double-stranded DNA molecules through semi-reserved replication, which is passed to the offspring by the parent methylation mode of another chain without methylation. Denovo methylation is a type of DNA methylation that occurs when different DNA MTases catalyze two strands of DNA without methylation [16]. C5-Mtases in plants fall into four categories, MTase (MET), chromomethylase (CMT), domains rearranged MTase (DRM), and Dnmt2 [17, 18]. MET is mainly used in methylation of the heterochromatin region of the CG site of the symmetric sequence, which is a very important part of the methyltransferase [19]. CMT is a specific type of DNA MTases that maintains CHG and CHH site methylation and, to a certain extent plays a role in stabilizing the heterochromatin state of the genome [20]. The function of DRM is to catalyze the methylation of cytosine and to maintain the cytosine methylation of non-CpG sites under the guidance of RNA [21]. DRM is homologous to the DNMT3 of animals [22, 23]. The proteins encoded by the plant Dnmt2 family are very similar to those of mice, bacteria and yeast C5-Mtases, but their mechanism of action in the process of C5 methylation has not been clarified [18].

Cytosine-5 DNA MTases were discovered in 1925 by Robert D. Coghill [24]. DNA methylation catalyzed by MTases was observed in bovine thymus in 1948 by Hotchkiss [25]. In 1964, Gold and Hurwitz identified the first DNA MTases in *Escherichia coli* [26]. Besto identified the first plant DNA MTases, and MET was isolated from *Arabidopsis thaliana*. Its encoded protein, AtMET1, had high homology with the methylation enzyme Dnmt1 in mice [27]. The reaction mechanism of DNA methylation was first explained in prokaryotic organisms in 1993 by Finnegan [28]. DNA MTases were identified in *Arabidopsis thaliana*, rice, maize, solanaceae, Tobacco, Legumes and other plants [22, 29-32].

## Materials and methods

### Identification of cotton DNA MTases family members

The cotton genome information was downloaded from CottonGen (https://www.cottongen.org/), and the DNA-methylase structure domain (PF00145) was downloaded from the Pfam (http://pfam.xfam.org/) database (IPR001525). The DNA-methylase.hmm Hidden Markov model was constructed with DNA-methylase.hmm as the reference HMMER3.0 (http://hmmer.org/download.html). The cotton genome database was queried to obtain the gene location and name of candidate protein family members containing DNA-methylase structure domains in cotton and to obtain GFF (general feature format) files from genome annotation files. Then, we obtained the gene position on the chromosome and used local BLAST 2.2.31+ to obtain the CDS sequence and protein sequence of the corresponding gene and obtained the whole sequence of the gene corresponding to the genome based on its position on the chromosome. The gene protein sequence is in the Smart software. The Pfam30.0 database is analyzed to ensure that each candidate gene contains a DNA-methylase structure domain. Subcellular location prediction was performed on cello. The ProtParam was obtained by protein analysis.

### Cotton DNA MTases gene structure, and evolutionarily conserved protein domain analysis

The CDS sequence of cotton DNA MTase genes and the corresponding genome-wide sequence in GSDS2.0 (http://gsds.cbi.pku.edu.cn/) were used to map the gene structure. The software Smart (http://smart.embl-heidelberg.de/) was used to determine the protein conserved structural domains.

### Phylogenetic analysis

Cotton DNA-methylase (PF00145, IPR001525) was used as the key word in Phytozome v12.1 (https://phytozome.jgi.doe.gov/pz/portal.html) in the database rather than the homologous sequences of other species (file 2). Clustal W software was used to analyze the amino acid sequence alignment. MEGA7.0 software was used to construct the phylogenetic tree with the neighbor-joining method. The number of bootstraps was 1000.

### Expression pattern analysis of cotton DNA MTases under stresses

The phytotron sand culture cultivation method was used for *G*.*raimondii* and *G. arboreum* L. *Shixiya* 1 (16h light / 8h dark, day 28°C, night 25°C). *G. hirsutum* L. TM-1 processing salt (200mM) and PEG6000 (20%) were used at the three-leaf stage at 0h, 1h, 3h, 6h, and 12h. Total RNA was extracted from root, stem, and leaf samples and reverse transcribed into cDNA. The primers for the real-time fluorogenic quantitative PCR were designed with the NCBI-line primer design tool primer-BLAST (http://www.ncbi.nlm.nih.gov/tools/primer-blast/) (file 1). The RNA was reverse transcribed to cDNA samples as a template for quantitative PCR experiments using *G*.*raimondii, G. arboreum* L. *Shixiya* 1, and *G. hirsutum* L. TM-1 to determine the gene expression of DNA MTases. The reaction conditions were 94°C for 30s; 40 cycles of 94°C denaturation for 5s, 50°C annealing for 34s, extension at 72°C for 10s;, and a final extension for 34s at 72°C.

### PYL156:VIGS silencing and Relative expression of GhDMT6 gene

The plasmid pYL156 was digested by EcoR and Xma I, and the digested product was detected by 1.2% agarose gel. The carrier segment was recycled. In-Fusion technique was used to insert t he VIGS silencing fragment into the vector pYL156, and then transferred into DH5a. The positive clones were selected. PCR detection and sample sequencing were carried out to obtain the correct monoclonal. The vector pYL156:*GhDMT6* was successfully constructed.

TM-1(*Gossypium hirsutum* L.) was cultured in the artificial climate chamber by sand culture method. After about 5 days of seedling emergence, the preserved Agrobacterium tumefaciens wer e resuscitated and transferred to 60 ml LB liquid medium. The cultured Agrobacterium tumefaciens were shaken to OD600=1.5, centrifuged for 5 minutes with 5 000 rpm of bacterial liquid, and the supernatant was discarded. Using 45ml(10mM MES+10 mMgCl2+200 AS) resuspension to culture the thallus. In order to remove a small amount of antibiotics, repeat the operation, then stewing the thallus in 25°C for 4h.

After mixing the suspension solution containing pYL156:*GhDMT6*, pYL156 and pYL156:C LA1 with the suspension solution containing auxiliary carrier pYL192 isopyknic, the cotton was prepared to be infected. A little pore was stabbed at the back of cotyledon with a sterile needle, and suspension solution injected into the cotyledon spread over the whole cotyledon. After inoculation, the cotton seedlings were put back into the artificial climate chamber for dark culture at 23°C for 24h, and then cultured at 23°C with other conditions unchanged. When the albinism symptoms of the positive control seedlings were obvious, the cotton without infection and pYL156:GhDMT6 were treated with 200 mM NaCl4°C.

## Results

### Genome-wide identification of cotton DNA MTases family members

A total of 51 DNA MTase members were identified from the whole genome of cotton. Group A had 8 DNA MTases and group D had 9 DNA MTases, which were named *GaDMT1*- *GaDMT8* and *GrDMT1*- *GrDMT9*, respectively, according to their sequence on the chromosomes. Similarly, 16 DNA MTases were identified in the AD_1_ group, named *GhDMT1*- *GhDMT16*, and 18 DNA MTases were identified in the AD_2_ group, named *GbDMT1*- *GbDMT18*. Most of the DNA MTases in the four cotton species are located on the chromosome. There are 161-1577 different amino acids, and most contain 300-800 amino acid residues. Because of the difference in the regional gene structure of the N-terminal, *GbDMT3* was up to 1577 amino acids, whereas *GbDMT17* contains only 161 amino acids. The structure of the gene is composed of 118-402 different amino acids, and the theoretical pi (PI) ranges from 4.67 to 9.24. The predicted subcellular localization shows that most DNA MTases are located in the cytoplasm, but some are located in the outer membrane. *GbDMT13* is predicted to be periplasmic (table 1).

**Table 1.** Basic characteristics of DNA Mtase genes in the cotton genome

| gene name | Accession number | Location (chromosome) | Position (domain) | CDS (bp) | AA | PI | Predicted subcellular localization |
| --- | --- | --- | --- | --- | --- | --- | --- |
| GaDMT1 |  | CA_chr1: 66883217:66889872- | 499-871 | 2724 | 907 | 5.09 | Cytoplasmic |
| GaDMT2 | Cotton_A_27044 | CA_chr2: 55459962..55465882- | 480-771 | 4638 | 1545 | 5.66 | Cytoplasmic |
| GaDMT3 | Cotton_A_20175 | CA_chr3: 24408653..24412255+ | 13-389 | 1179 | 637 | 6.02 | OuterMembrane |
| GaDMT4 | Cotton_A_29334 | CA_chr3: 96706846..96713328- | 552-692 | 2082 | 693 | 7.44 | OuterMembrane |
| GaDMT5 | Cotton_A_29333 | CA_chr3: 96716269:96725942- | 475-822 | 2580 | 859 | 6.24 | Cytoplasmic |
| GaDMT6 | Cotton_A_36034 | CA_chr10: 113686352..113690340+ | 513-631 | 1914 | 663 | 4.8 | Cytoplasmic |
| GaDMT7 | Cotton_A_08447 | CA_chr12: 1201266..1207466- | 534-653 | 1992 | 392 | 4.94 | Cytoplasmic |
| GaDMT8 | Cotton_A_27234 | CA_chr12: 56291689..56297548- | 1110-1539 | 1494 | 497 | 5.82 | Cytoplasmic |
| GaDMT9 | Cotton_A_19737 | CA_chr13: 63213811..63219313+ | 155-372 | 2421 | 806 | 5.25 | Cytoplasmic |
| GrDMT1 | Cotton_D_gene_10004803 | Chr2:8382256..8388371 + | 494-785 | 2463 | 820 | 5.67 | Cytoplasmic |
| GrDMT2 | Cotton_D_gene_10010304 | Chr3:6796286..6800300- | 512-630 | 1911 | 636 | 4.8 | Cytoplasmic |
| GrDMT3 | Cotton_D_gene_10010121 | Chr4:457766..461539 + | 13-389 | 1203 | 400 | 5.55 | OuterMembrane |
| GrDMT4 | Cotton_D_gene_10027875 | Chr4:20122791..20126504- | 511-630 | 1923 | 640 | 4.9 | Cytoplasmic |
| GrDMT5 | Cotton_D_gene_10009363 | scaffold141: 194013:200650- | 495-869 | 2718 | 905 | 5.1 | Cytoplasmic |
| GrDMT6 | Cotton_D_gene_10000349 | scaffold531:66538..72372 - | 1128-1557 | 4692 | 1563 | 5.51 | Cytoplasmic |
| GrDMT7 | Cotton_D_gene_10002270 | scaffold372: 306669:314046- | 530-928 | 2892 | 963 | 7.02 | OuterMembrane |
| GrDMT8 | Cotton_D_gene_10002271 | scaffold372: 317040:326457- | 789-1165 | 3609 | 1202 | 8.44 | OuterMembrane |
| GhDMT1 | CotAD_37635 | At_chr6:11311066..11315214- | 512-630 | 1911 | 636 | 4.8 | Cytoplasmic |
| GhDMT2 | CotAD_51709 | At_chr9:63179674..63185543- | 1076-1505 | 4536 | 1511 | 6.34 | Cytoplasmic |
| GhDMT3 | CotAD_46796 | At_chr9:65748738..65752451+ | 511-630 | 1923 | 640 | 4.85 | Cytoplasmic |
| GhDMT4 | CotAD_10542 | Dt_chr1:40856014..40859213- | 1-308 | 1035 | 344 | 7.76 | Cytoplasmic |
| GhDMT5 | CotAD_49037 | Dt_chr2:8190450..8196335- | 496-781 | 2451 | 816 | 5.62 | Cytoplasmic |
| GhDMT6 | CotAD_04205 | Dt_chr5:13787674..13791315+ | 13-389 | 1206 | 401 | 5.82 | OuterMembrane |
| GhDMT7 | CotAD_13275 | Dt_chr6:40651384..40655909- | 512-611 | 1878 | 625 | 4.72 | OuterMembrane |
| GhDMT8 | CotAD_24264 | Dt_chr7:37416068..37421133+ | 230-632 | 2007 | 668 | 4.81 | Cytoplasmic |
| GhDMT9 | CotAD_00990 | Dt_chr10: 6477855..6487242+ | 789-1157 | 3585 | 1194 | 8.51 | OuterMembrane |
| GhDMT10 | CotAD_00992 | Dt_chr10:6495924..6502242+ | 299-680 | 2094 | 697 | 6.75 | Cytoplasmic |
| GhDMT11 | CotAD_14980 | scaffold39.1:602870..611908+ | 667-846 | 2646 | 881 | 6.94 | OuterMembrane |
| GhDMT12 | CotAD_18652 | scaffold71.1:1581335..1591239- | 547-666 | 2031 | 676 | 4.91 | Cytoplasmic |
| GhDMT13 | CotAD_41398 | scaffold294.1:1053646..1057162- | 319-500 | 1533 | 510 | 5.45 | Cytoplasmic |
| GhDMT14 | CotAD_41399 | scaffold294.1:1063401:1074111- | 780-1161 | 3597 | 1198 | 8.71 | OuterMembrane |
| GhDMT15 | CotAD_46012 | scaffold1041.1:189137..194922+ | 1112-1541 | 4644 | 1547 | 5.57 | Cytoplasmic |
| GhDMT16 | CotAD_40093 | scaffold2005.1:29096..35764+ | 453-828 | 2595 | 864 | 5.02 | Cytoplasmic |
| GbDMT1 | Gbscaffold4563.2.0 | At01: 75311449..75320269 | 441-816 | 2556 | 851 | 5.66 | Cytoplasmic |
| GbDMT2 | Gbscaffold7611.1.0 | At05:80836461..80840887+ | 564-683 | 2082 | 693 | 5.03 | Cytoplasmic |
| GbDMT3 | Gbscaffold10176.3.0 | At05:85393045..85400787+ | 1142-1571 | 4734 | 1577 | 5.49 | Cytoplasmic |
| GbDMT4 | Gbscaffold23728.21.0 | At07: 5162033..5169156 | 503-875 | 2739 | 912 | 4.98 | Cytoplasmic |
| GbDMT5 | Gbscaffold11439.6.0 | At08:92243382..92245846- | 278-428 | 1431 | 476 | 5.77 | Cytoplasmic |
| GbDMT6 | Gbscaffold11439.5.0 | At08: 92252927..92259967 | 131-508 | 1641 | 546 | 8.13 | Cytoplasmic |
| GbDMT7 | Gbscaffold10104.36.0 | At08:106991133..106995128+ | 13-389 | 1179 | 392 | 5.4 | OuterMembrane |
| GbDMT8 | Gbscaffold155.2.0 | At09:9281821..9292014+ | 1068-1186 | 3579 | 1192 | 4.78 | Cytoplasmic |
| GbDMT9 | Gbscaffold155.3.0 | At09:9298124..9301305+ | 374-492 | 1497 | 498 | 5.56 | Cytoplasmic |
| GbDMT10 | Gbscaffold10257.5.0 | Dt01: 55556607..55564136 | 438-672 | 2370 | 789 | 5.78 | Cytoplasmic |
| GbDMT11 | Gbscaffold9562.4.0 | Dt04:6380024..6388975+ | 1126-1555 | 4686 | 1561 | 5.6 | Cytoplasmic |
| GbDMT12 | Gbscaffold9562.5.0 | Dt04:6390322..6394029+ | 475-834 | 2523 | 840 | 6.49 | Cytoplasmic |
| GbDMT13 | Gbscaffold12826.2.0 | Dt04:9853138..9856849+ | 511-630 | 1923 | 640 | 4.93 | Cytoplasmic |
| GbDMT14 | Gbscaffold12826.3.0 | Dt04:9869411..9871104+ | 219-338 | 1047 | 348 | 8.6 | Cytoplasmic |
| GbDMT15 | Gbscaffold8153.13.0 | Dt07: 5581094..5588192 | 492-866 | 2709 | 902 | 5.2 | Cytoplasmic |
| GbDMT16 | Gbscaffold34888.4.0 | Dt08:67685650..67693483- | 19-642 | 1962 | 653 | 5.21 | OuterMembrane |
| GbDMT17 | Gbscaffold8535.6.0 | scaffold8535:88760..90489+ | 7-125 | 486 | 161 | 9.24 | Periplasmic |
| GbDMT18 | Gbscaffold12265.1.0 | scaffold12265:8693..15154+ | 576-694 | 2103 | 700 | 4.67 | Cytoplasmic |

### Multi-sequence alignment and evolutionary analysis

The DNA MTases in cotton have a C-terminal MTase catalytic region structure domain and a specific N-terminal domain, which is consistent with those of Arabidopsis, rice, maize, solanaceae, Tobacco, Legumes and other crops [22, 29-32]. To evaluate the evolution of DNA MTases in A, D, AD_1_ and AD_2_, multiple sequence alignment was performed and a phylogenetic tree was constructed for 51 members of the DNA MTase family (Fig. 1A). The DNA MTases in cotton are divided into four subfamilies, namely, CMT, MET, DRM, and Dnmt2 [33]. Among these 51 DNA MTases, the CMT subfamily contains 25 members, with 4, 5, 9, and 7 members in the D, A, AD_1_, and AD_2_ groups, respectively. There are three different types of CMTs: CMTa has 9 members, with 1, 2, 3, and 3 members in the D, A, AD_1_, and AD_2_ groups, respectively; CMTb has 6 members, with 1, 1, 2, and 2 members in the D, A, AD_1_, and AD_2_ groups, respectively; and CMTc has 10 members, with 2, 2, 4, and 2 members in the D, A AD_1_, and AD_2_ groups, respectively. MET has 7 members in the D, A, AD_1_, and AD_2_ groups. DRM has 14 members, with 2, 2, 4, and 6 members in the D, A, AD_1_, and AD_2_ groups, respectively. There are 5 members in Dnmt2, with 1, 1, 1, and 2 members in the D, A, AD_1_, and AD_2_ groups, respectively.

**Fig. 1.**
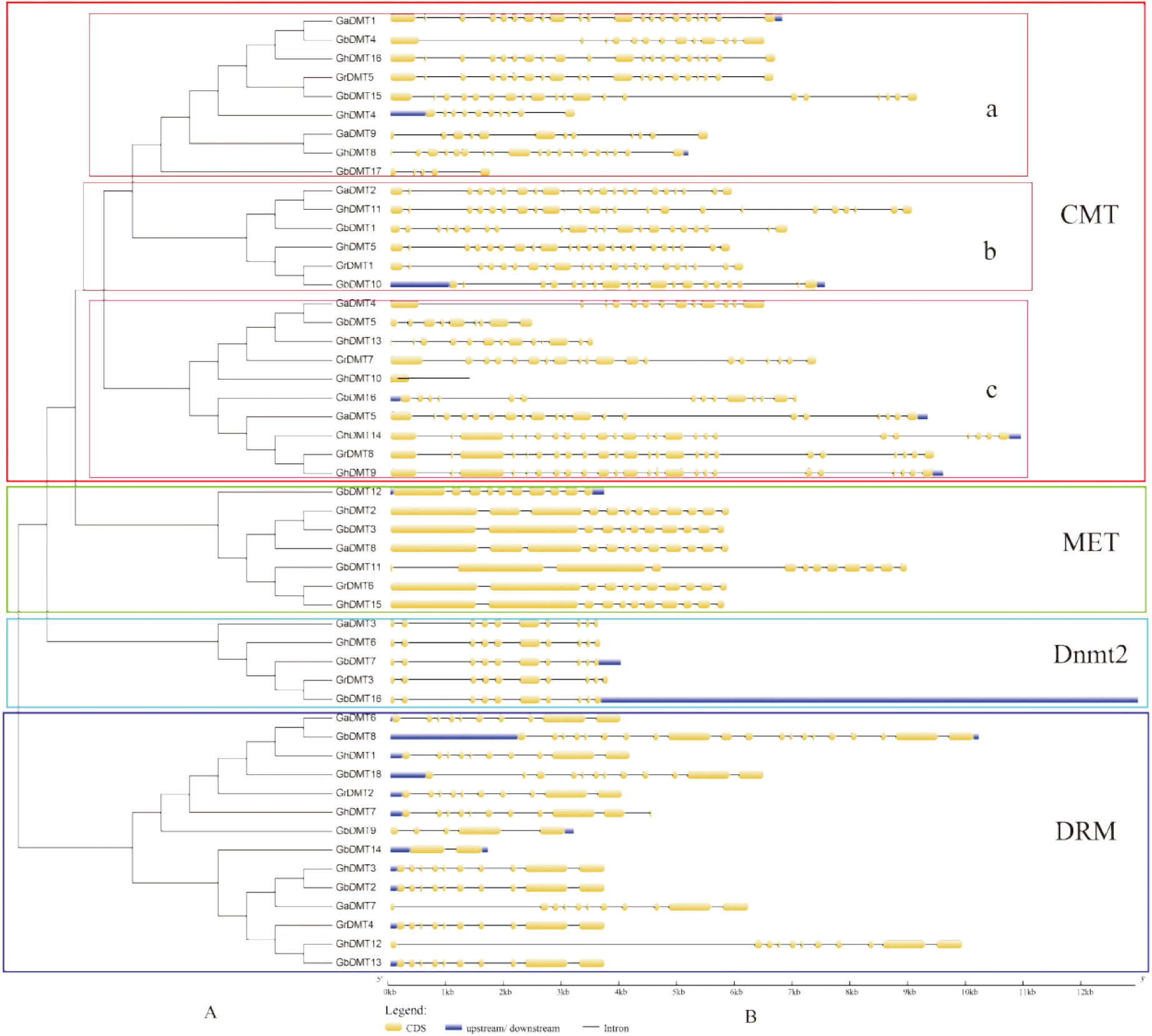
Phylogenetic analysis of the DNA Mtase gene family in cotton (A) and genetic structure (B). Cotton DNA Mtase family members are divided into four subfamilies, CMT, MET, DRM, Dnmt2.

### Genetic structure and protein domains of DNA MTases in cotton

Gene structure analysis is an important method in the study of genetic evolution. The number of introns and exons in DNA MTase family members in groups D, A, AD_1_ and AD_2_ were analyzed, and a DNA MTase gene structure for cotton was created (Fig. 1B). The results showed that the numbers of exons in different MTase genes in cotton were very different; the *GbDMT14* gene had the fewest exons, with only 2 exons, whereas *GrDMT8, GhDMT9, GhDMT11* and *GhDMT14* had 24 exons. The Dnmt2 family contains only 10 exons, the Met family contains 10-12 exons, the CMTa family contains 5-22 exons, the CMTb family contains 19-24 exons, the CMTc family contains 9-24 exons, and the DRM family contains only 2-20 exons.

Motif analysis of the 51 DNA MTase proteins in cotton is shown in Fig. 2 The C-terminal catalytic region has 6 highly conserved motifs: motif x and motif i for Sam-binding sites; motif iv, motif vi, motif viii, and motif ix are the c5-Mtase functional catalytic sites, of which motif iv is the active site, motif vi is the target cytosine binding site. motif viii is the DNA neutralization region, and motif ix is the target sequence location identification area, which is consistent with the related literature [22]. The MTases of cotton DNA have different motif orders. The motif order of the DRM family is X, VI, VIII, IX, I, and IV, and the order for the DNMT2 family is I, x, vi, viii, iv, and ix. There are three motif orders in CMT: The first is I, iv, vi, x, viii, and ix; the second is IX, i, iv, vi, x, and viii; and the third is VI, i, iv, x, viii, and ix.

**Fig. 2.**
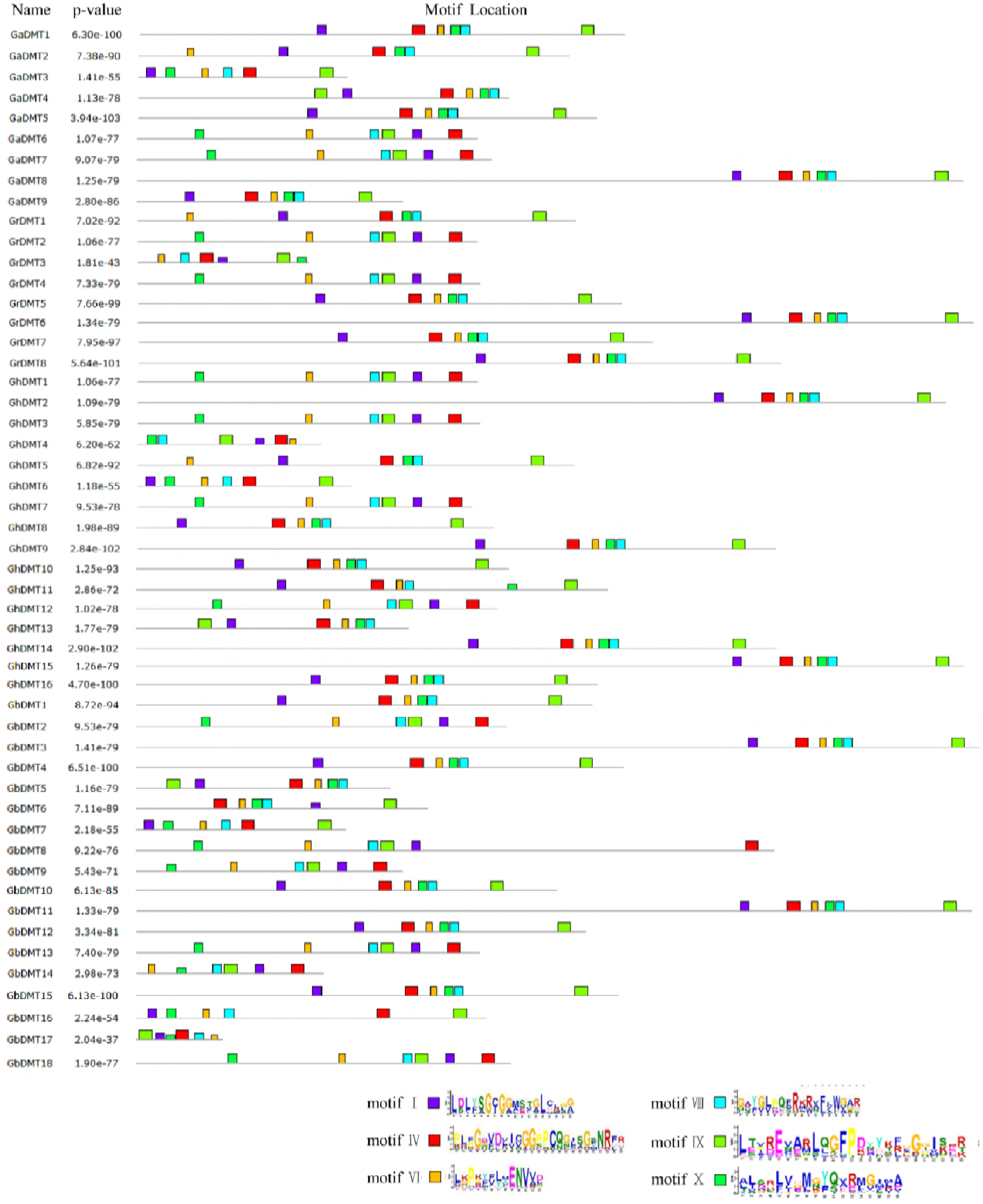
Analysis of the distribution of cotton DNA Mtase protein motifs using online software. The rectangular length conforms to the length of the motif. The order and position of the motif correspond to the order and position of the bases in a single protein sequence.

**Fig. 3.**
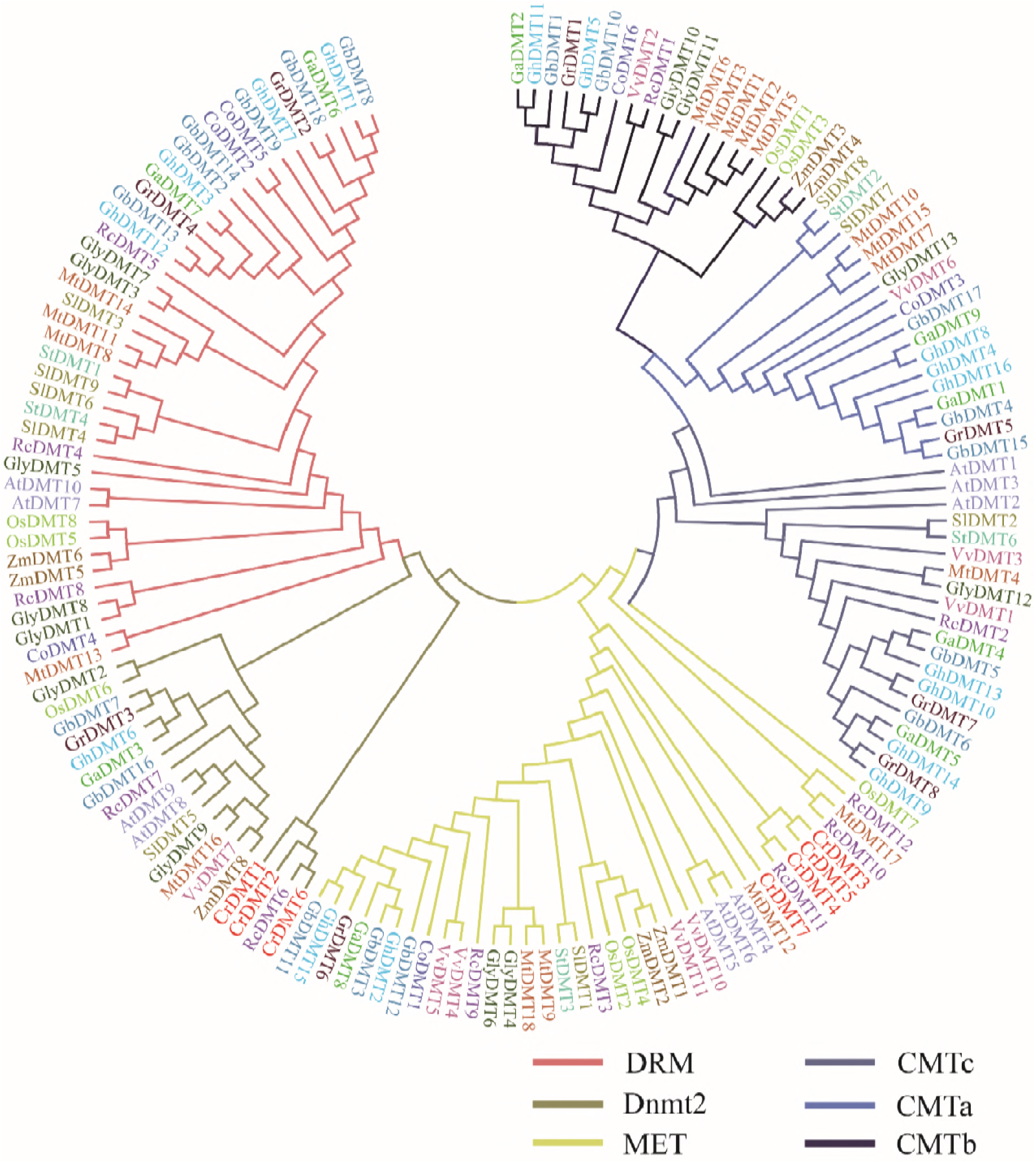
DNA Mtase family phylogenetic tree of cotton and other crops. The neighbor-joining method was used to construct the tree without a root system, and the value of the bootstrap was 1000.

### The relationship between the cotton DNA MTase family and DNA MTases in other crops

Phylogenetic trees are used to reveal the homologous and evolutionary relationships of DNA MTase families from different species. To show the evolutionary relationship between the members of the cotton DNA MTase family and those of Arabidopsis, cocoa, Medicago, rice, Chlamydomonas reinhardtii and other crops, the amino acid sequences of DNA MTase family members in several crops were compared (table 2) to the DNA of the MET, DRM, and CMT family members. In monocotyledons, the differentiation of DNA MTases was separate from the evolution of the dicotyledons, and the Dnmt2 family did not differentiate in the monocotyledons, which showed that the Dnmt2 family was highly conserved during evolution. The DNA MTases in cotton are closer to those in cocoa in various branches, suggesting that they have similar functions. (Note: The VvDMT8, VvDMT9, ZmDMT7, and StDMT5 sequences are shorter and cannot be used in the construction of the inter-species evolutionary tree.)

**Table 2.** Basic information of related species in analyzing the phylogenetic tree

| species name | short | classification | species name | short | classification |
| --- | --- | --- | --- | --- | --- |
| *Ricinus communis* | Rc | D | *Chlamydomonas reinhardtii* | Cr | A |
| *grapevine* | Vv | D | *Medicago truncatula* | Mt | D |
| *Oryza sativa* | Os | M | *Arabidopsis thaliana* | At | D |
| *Glycine max* | Gly | D | *Solanum lycopersicum* | Sl | D |
| *Zea mays* | Zm | M | *Solanum tuberosum* | St | D |
| *cacao* | Co | D |  |  |  |

D, M and A represent Dicotyledon, Monocotyledon and Algae, respectively.

### Cotton gene expression analysis of different DNA MTases under stress

To study the expression patterns of DNA MTase genes of cotton in different tissues under salt and drought stress, the *G*.*raimondii, G. arboreum* L. *Shixiya* 1, and *G. hirsutum* L. TM-1 were developed to the trefoil stage, and real-time quantitative PCR was performed. The results showed that the three cotton species had different expression patterns under different stress conditions, and *G*.*raimondii, G. arboreum* L. *Shixiya* 1, and *G. hirsutum* L. TM-1 had obvious tissue differences when treated with PEG6000 and NaCl. *G. arboreum* L. *Shixiya* 1 expressed more genes in the cut root and stem under salt and drought stress. The leaf also had more genes, and with different treatments, the gene expression was also different. *GaDMT7* and *GaDMT4* decreased in the stem under salt treatment, and drought treatment also decreased their expression. *G*.*raimondii* also has organizational specificity. Gene expression was decreased in the rhizome and increased in the leaf. *GrDMT5* decreases in the stem but increases in the root performance under the salt treatment, showing that the same gene is expressed differently in various tissues. *GrDMT1* is decreased in leaves under drought treatment; however, salt treatment increases its expression. For *G. hirsutum* L. TM-1 and the above two cotton varieties, the genes are mainly distributed in the root of the rhizome. Additionally, the leaf has more genes. *GhDMT9* is expressed in the root under salt treatment. In the leaf, expression first increases and then decreases.

### Functional analysis of GhDMT6 gene in cotton

After 6 days of natural drought, the plant phenotypes were significantly different. The plants injected with pYL156 and those of the wild type were yellow and withered, seriously dehydrated, and the plants died. The plants injected with pYL156:*GhDMT6* had no true leaf blight and water loss (Fig. 5a). The relative expression level of *GhDMT6* gene in pYL156:*GhDMT6* infected cotton seedlings was detected. The figure shows that the expression level of *GhDMT6* gene in pYL156 infected cotton seedlings did not change under different stresses. Compared with the control, the transcription level of *GhDMT6* gene in pYL156 infected cotton seedlings decreased the most in the stem, followed by the leaves, and decreased the least in the roots (Fig. 5d).

**Fig. 4.**
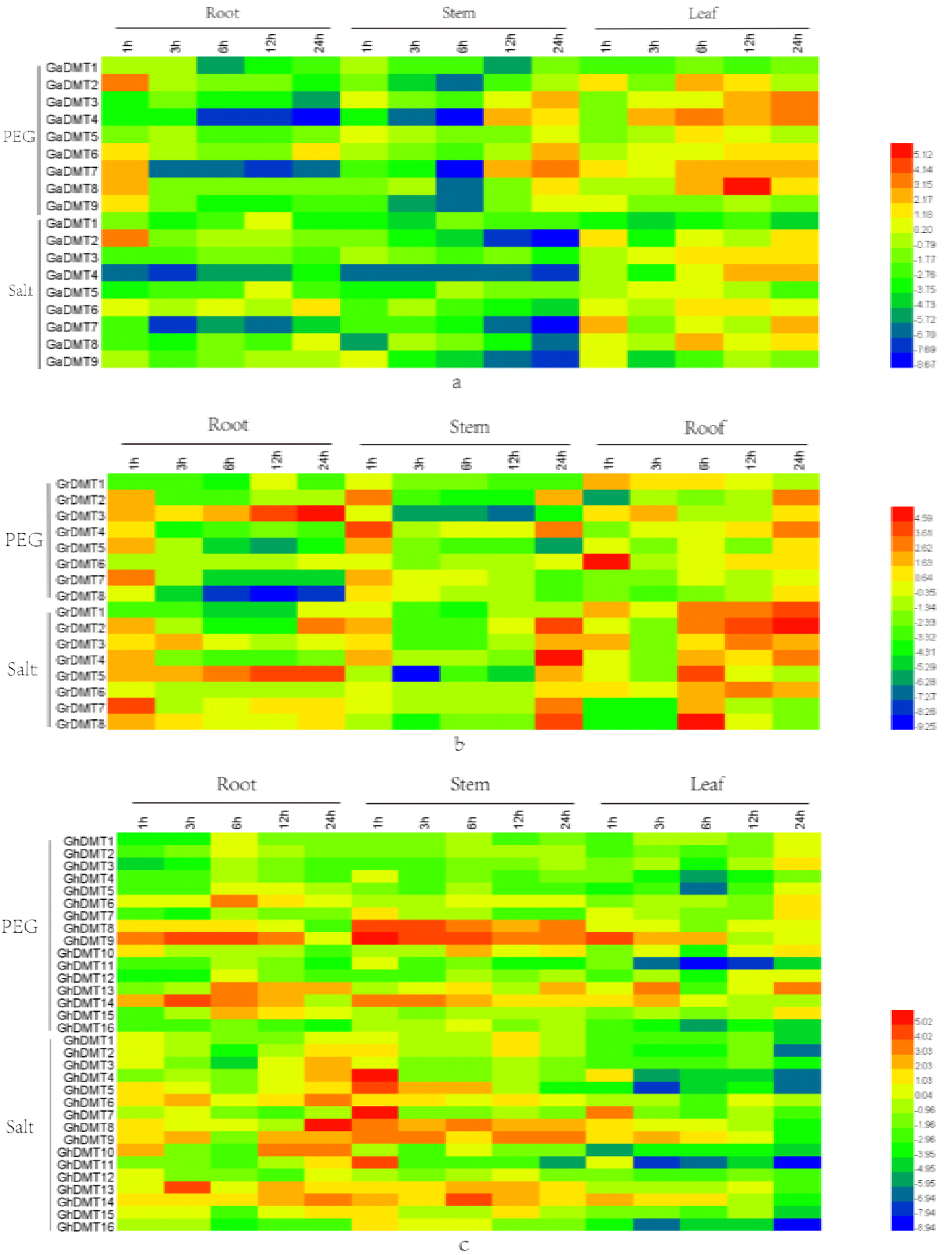
The expression of DNA Mtase genes in the roots, stems and leaves of three cotton species, namely, *Shixiya 1,G. raimondii*, and *G. hirsutum* L. TM-1, under drought and salt treatments at different times. a *G. arboreum* L. *Shixiya* 1, b *G*.*raimondii*, and c *G. hirsutum* L. TM-1.

**Fig. 5.**
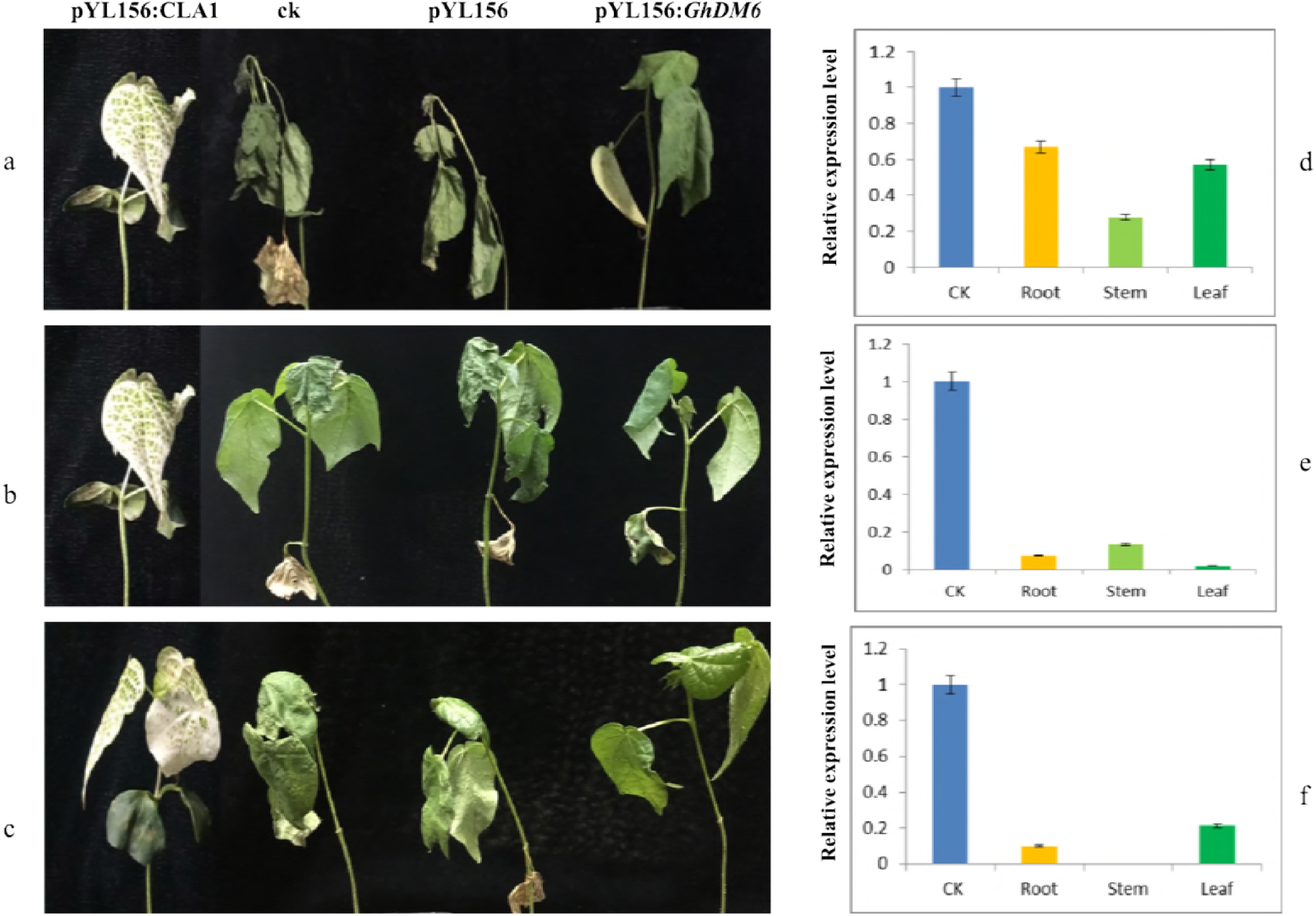
Relative expression of GhDMT6 gene in different tissues of differen stress in Gossypium. hirsutum. a. Phenotypic differences of cotton after drought stress. b. Relative expression of GhDMT6 gene after drought stress in G. hirsutum L. TM-1. c. Phenotypic differences of cotton after cold stress. d. Relative expression of GhDMT6 gene after cold stress in G. hirsutum L. TM-1. e. Phenotypic differences of cotton after salt stress. f. Relative expression of GhDMT6 gene after salt stress in G. hirsutum L. TM-1

About 15 days after VIGS infection, the positive control plants showed obvious bleaching and other indexes were normal. Using no infection and pYL156 infection of cotton seedlings as control, the silent plants were treated with cold(4°C), drought (natural drought), salt(200 mM Nacl). After 36h of cold treatment, the phenotypic differences were obvious (Fig. 5b). The plants injected with pYL156 and wild-type true leaf wilted and curly, and the plants injected with pYL156:*GhDMT6* gene grew normally without phenotypic changes. The expression level of *GhDMT6* gene in pYL156 infected cotton did not change under different stresses, and the transcriptional level of *GhDMT6* gene in pYL156:*GhDMT6* infected cotton was significantly reduced compared with the control. Leaf blade decreased the most, followed by root, stem the least (Fig. 5e).

The phenotype of cotton seedlings was significantly different after 3 days, which were treated with 200mM NaCl. The seedlings of pYL156 and wild-type cotyledons were exfoliated, the leaf edges of true leaves were severely coked, and the plants with pYL156:*GhDMT6* gene were not withered and dehydrated (Fig. 5c). The results showed that the expression of *GhDMT6* gene in cotton seedlings infected with pYL156:*GhDMT6* did not change under salt stress. The transcription level of *GhDMT6* gene in cotton seedlings infected with pYL156:*GhDMT6* decreased most in stems, followed by roots, and decreased least in leaves (Fig. 5f).

## Discussion

With the completion of the genome project of cotton [34], the identification and study of gene family classifications, evolutionary features and function prediction at the whole-genome level is a hotspot of cotton functional gene research. Cotton is one of the pioneer plants in saline-alkali lands. DNA MTases are key enzymes in DNA methylation, which is closely related to resistance to stress. Therefore, the study of genome-wide DNA MTases is of great significance to cotton breeding, the identification of functional genes and the mechanism of cotton resistance.

DNA methylation affects many biological processes, including disease-associated syndromes in humans [35]. Natural variations of epialleles play a role in plant evolution [36], morphological diversity in plants [37], and the selection and breeding of agronomic traits in crops [4, 38]. Increasing evidence from recent studies suggests that DNA methylation plays an important role in regulating the stress response/adaptation in plants. DNA methylation may be an adaptation mechanism of plants in response to adversity. Osmotic stress causes DNA methylation in the chromatin region of tobacco (*Nicotiana* L.) and tissue culture cells [39]. Salt stress can inhibit *zmPP2C* expression and induce *zmGST* expression in maize (*Zea mays*) seedlings [40]. Heavy metals, such as Cd and Pb, can increase the methylation level in the genomes of rice, wheat, rapeseed and other crops and produce toxicity [41].

This study, for the first time, systematically analyzed the DNA MTase gene family in the cotton genome, and 51 DNA MTases were obtained, divided into four Asian families. There are 7 members of the Met family with a region rich in glutamic acid and aspartate, similar to those of *Arabidopsis thaliana*. Although the amino acid residues in this region are important for the function of DNA MTases, the specific effect has not been determined [42]. The CMT family has 25 members, and the main sites are CHG sequences. They have particularly high abnormal chromatin content, possibly because CHG methylation maintains the plant genome regional chromatin state [43]. The DRM family, which has 14 members, contains a ubiquitin-related structural domain (UBA) and an interface between seat proteins, which introduces DRM to specific DNA regions to complete methylation of the region. The DNMT2 family has 5 members and is highly homologous in animals, plants, and prokaryotic organisms. This high homology may be due to the evolution of prokaryotic organisms. Its functional substrate is RNA, and the main target is tRNA^Asp^. DNMT2 family members can specifically methylate the tRNA^Asp^ of the reverse codon ring 38C [44]. There were 8 members in group A, 9 in group A, 16 in the AD_1_ group and 18 in the AD_2_ group. The results showed that there were more DNA MTase genes in the AD genome than in the A genome or D genome, but the number of genes in the AD genome was not equal to the sum of genes in genome A and genome D. The number of DNA MTases in the AD_1_ genome is less than the sum of genomes A and D, which may be associated with genetic loss during the evolution of the twofold ancestral AD_1_ genome. The number of DNA MTases in the AD_2_ genome is greater than the sum of genomes A and D, which may be related to gene duplication during the evolution of the twofold ancestral AD_1_ genome. However, the quantitative difference between the DA_1_ genome and the DA_2_ genome may be the difference between natural selection and artificial domestication [45].

Gene expression profiling is usually related to gene functional identification. For the three cotton species under salt and drought stress, DNA MTase gene expression is significantly different in different tissues. This differential gene expression may be caused by the corresponding adverse effects of the regulatory pathway. This shows that different DNA methylation enzymes have different functions and participate in many regulatory pathways and that the plant’s response to adversity is the result of synergistic effects through multiple pathways and is a complex process. Drought and salt stress can cause osmotic pressure, and the change in the DNA MTase gene transcription level in cotton may be caused by the joint action of these two types of stress. This is similar to the findings for other crops. Different cotton species in the same subfamily have different patterns of expression, and DNA MTase gene may have specific functions.

The function of the GhDMT6 gene was verified by the VIGS silence. After silencing the cotton GhDMT6 gene, it was less sensitive to cold, salt and drought stress than the control group, which indicated that the gene had a certain significance in improving cotton resistance. In summary, the GhDMT6 gene is a gene involved in stress resistance.

### Funding information

This study was funded by the National Key Research and Development Program of China (2018YFD0100401).

### Compliance with ethical standards

The experiments described here comply with the current laws of China.

